# FGF8-mediated signaling regulates tooth developmental pace during odontogenesis

**DOI:** 10.1101/2020.09.16.299388

**Authors:** Chensheng Lin, Ningsheng Ruan, Linjun Li, Yibin Chen, Xiaoxiao Hu, YiPing Chen, Xuefeng Hu, Yanding Zhang

## Abstract

The developing human and mouse teeth constitute an ideal model system to study the regulatory mechanism underlying organ growth control due to the fact that their teeth share highly conserved and well-characterized developmental processes and their developmental tempo vary notably. In the current study, we manipulated heterogenous recombination between human and mouse dental tissues and demonstrate that the dental mesenchyme dominates the tooth developmental tempo and FGF8 could be a critical player during this developmental process. Forced activation of FGF8 signaling in the dental mesenchyme of mice promoted cell proliferation, prevented cell apoptosis via p38 and perhaps PI3K-Akt intracellular signaling, and impelled the transition of the cell cycle from G1- to S-phase in the tooth germ, resulting in the slowdown of the tooth developmental pace. Our results provide compelling evidence that extrinsic signals can profoundly affect tooth developmental tempo and the dental mesenchymal FGF8 could be a pivotal factor in controlling developmental pace in a non-cell-autonomous manner during mammalian odontogenesis.

## Introduction

Although mammals possess an almost identical body plan with morphologically and functionally similar organs and systems, their developmental programs and body sizes vary greatly among different species. The intrinsic developmental programs, together with extracellular signals, control the individual organ size and thus body size ^(1)^. Generally, larger-bodied species own longer gestation lengths with slower developmental tempos to acquire larger body sizes ^(2,3)^. Obviously, developmental tempo and organ size are two closely interrelated variables during individual growth (i.e. embryonic development and postnatal growth) and morphogenesis, both of which must be tightly harmonized to ensure the correct establishment of body plan and therefore are the result of a precisely controlled process.

The mechanisms that regulate organ size have been well investigated in non-mammalian models, particularly in *Drosophila melanogaster*, in which roles of TOR (target of rapamycin), insulin/IGF (insulin-like growth factor), Ras/Raf/MAPK, JNK (c-Jun N-terminal kinase), and Hippo signaling pathways are well defined ^(4)^. In mammals, Hippo and IGF pathways have been evidenced to be involved in controlling organ size ^(5-10)^. Additionally, several conserved families of secreted growth factors, including FGF (fibroblast growth factor), BMP (bone morphogenetic protein), and TGF-β (transforming growth factor-β), are deemed to be involved in the regulation of individual growth due to their ability to modulate cell proliferation and apoptosis during embryonic development ^(4,11,12)^. However, the regulatory mechanisms of developmental pace in mammals are remain largely unknown.

Mammalian teeth have long served as a model organ for studying fundamental questions in organ development on the molecular basis ^(13)^. The main features of tooth morphogenesis are conserved in the mammalian lineage ^(14-16)^, as they are all formed by sequential and reciprocal interactions between epithelium and mesenchyme and share similar processes during tooth development ^(15)^. Despite these considerable homologies, mammalian dentition undergoes remarkable morphological diversification, displaying great variations not only in tooth shape, number, and size but also in developmental pace ^(13,15,17-19)^. Obviously, the period of tooth development in humans takes about 400 days from the initiation stage at embryonic 6^th^ week to tooth eruption at postnatal 6^th^ month. Whereas, this period in mice takes only about 20 days from the determination of tooth forming sites at E10.5 to tooth eruption at postnatal 10^th^ day ^(13)^. Thus, the human and mouse embryonic tooth provides an ideal model system for investigating the regulatory mechanism underlying the developmental tempo, however, the molecular mechanism that regulates the developmental pace of mammalian teeth remains elusive.

It is generally believed that diversified forms of organ morphogenesis (including the developmental tempo) among species are achieved by tinkering conserved signaling pathways instead of proposing novel ones during development ^(20-22)^. The difference in organismal growth rate and organ or body size between humans and mice primarily reflect the difference in cell number rather than the cell size ^(1)^. Many lines of evidence from mouse models demonstrate that numerous growth factors of conserved families, such as FGF, BMP, SHH, and Wnt, have been shown to play critical roles in tooth morphogenesis by regulating cell proliferation, differentiation, and apoptosis during odontogenesis ^(15,16,21,23-27)^, suggesting their potential roles in the regulation of the developmental tempo. Here, we provide compelling evidence that it is the dental mesenchyme that dominates tooth developmental tempo in a non-cell-autonomous manner and extrinsic FGF8 may act as a critical factor in controlling this process during odontogenesis.

## Materials and methods

### Mouse models

*Wnt1-Cre* and *R26R*^*Fgf8*^ mice were described previously ^(28,29)^. Dental mesenchyme-specifically overexpressed *Fgf8* mice were generated by crossing *Wnt1-Cre* mice with *R26R*^*Fgf8*^ mice. Mutant embryos were identified by PCR genotyping. All wild-type mice were on the CD-1 background. Mice were housed under the following identical conditions: 40-70% relative humidity, 12 hour light/dark cycle, and 20-25°C ambient temperature, in individually ventilated cages, with corncob granules for bedding and nesting, in groups of up to six. The mice were given free access to tap water and standard rodent chow (1010011; Jiangsu Xietong Pharmaceutical Bio-engineering Co., Ltd, Nanjing, China).

### Tissue recombination, organ culture, and subrenal culture

Fresh lower jaws isolated from human fetuses of 14^th^-16^th^ week gestation following medical termination of pregnancy were provided by Maternal and Child Health Hospital of Fujian Province, China. The precise embryonic age of fetuses was defined by the measurement of the crown-rump length (CRL) ^(30)^. Human and E13.5 mouse embryonic tooth germs were dissected out in ice-cold PBS and then treated with Dispase II at 37°C for 20 min. The dental mesenchyme and dental epithelium were separated with the aid of fine forceps and incubated on ice for further tissue recombination experiments. Three types of tissue recombination were conducted and are summarized in Table 1: E13.5 mouse dental epithelium with bell-stage human dental mesenchyme (14-16 weeks of gestation) (referred to as mde+hdm); E13.5 mouse dental mesenchyme with bell-stage human dental epithelium (hde+mdm); and E13.5 mouse dental mesenchyme with mouse dental epithelium from the same stage as control (mde+mdm). All recombinants were cultured in the Trowell-type organ culture dish overnight, followed by subrenal grafting in adult nude mice, as described previously ^(30,31)^. The use of animals and human tissues in this study was approved by the Ethics Committee of Fujian Normal University (permission #2011005).

**Table. 1.**
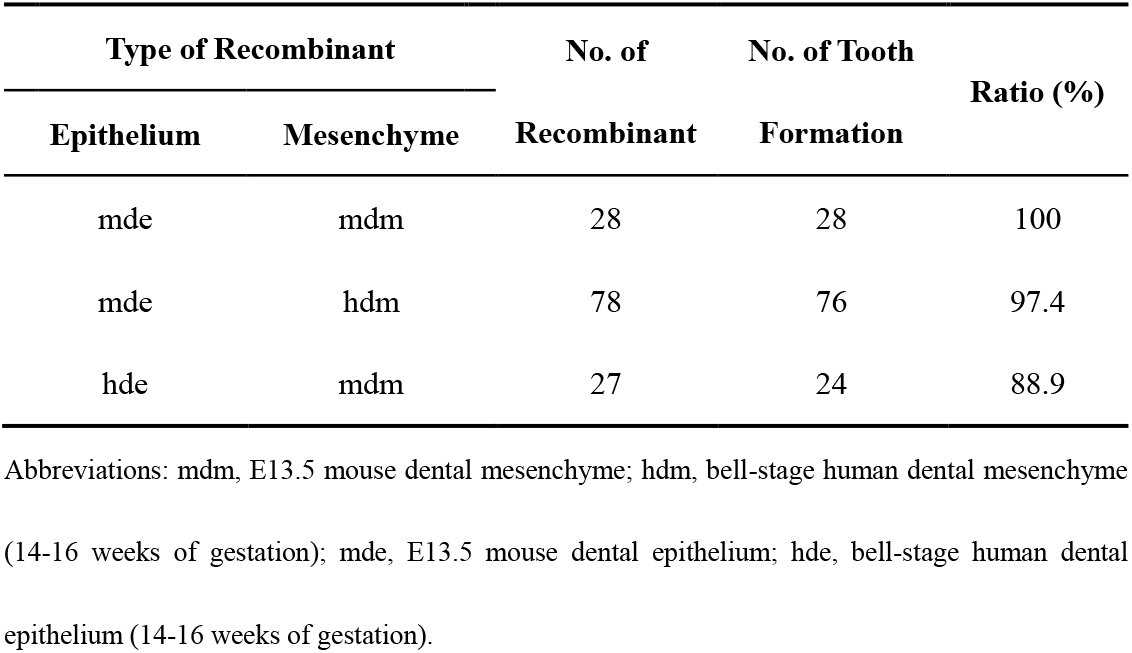
Ratio of tooth formation in different types of tissue recombinants.

### SU5402 injection

Two first mandibular premolar germs dissected out from the lower jaw of a human fetus of 16^th^ week gestation were cultured under the kidney capsules of two nude mice, one germ each. SU5402 (Calbiochem, California, USA) and DMSO (Merck, Darmstadt, Germany) was intraperitoneally injected into each recipient mouse at a dose of 25mg/Kg/d in 200 μL PBS twice a day for one week. The human tooth grafts were harvested for histological analysis after another week in subrenal culture.

### Lentivirus infection and cell re-aggregation

For efficient infection of lentivirus ^(32)^, isolated E13.5 mouse dental mesenchyme was digested with 0.25% trypsin at 37 °C for 5 min and dispersed into a single-cell suspension with the aid of a micropipette. To make a dental mesenchyme pellet, about 5×10^5^ dental mesenchymal cells were infected with 20 μL concentrated lentivirus in 200 μL culture medium in an Eppendorf tube. The tube was incubated at 37° in a 5% CO_2_ environment for 2 hours with its cap open, centrifuged at 3000 rpm for 3 min, and then incubated again in the same environment for 2 hours. Cell pellets were removed from the tubes, recombined with E13.5 dental epithelium, and cultured in the Trowell-type organ culture dish overnight. Recombinants were subsequently subjected to subrenal culture in nude mice.

### Histology, *in situ* hybridization, immunostaining, and Western blot

For histological analysis, harvested samples were fixed in 4% paraformaldehyde (PFA)/PBS, dehydrated, embedded in paraffin, and sectioned at 7 μm. Standard Hematoxylin/Eosin (H&E) and Azan dichromic staining were utilized for histological analysis ^(33)^. Ossified tissues or grafts were demineralized with EDTA/PBS for a week prior to dehydration. For *in situ* hybridization, dental tissues from staged human embryos were dissected out in ice-cold PBS and fixed in 4% RNase-free PFA/PBS at 4°C overnight. Human cDNAs, including *FGF1* (MHS6278-202758608), *FGF2* (EHS1001-208062206), *FGF15/19* (MHS6278-202756089), *FGF18* (MHS6278-202830446), were purchased from Open Biosystems (Alabama, USA) and were subcloned to pBluescript-KS plus vector for riboprobe transcription. Whole-mount *in situ* hybridization and section *in situ* hybridization were performed following standard protocols as described previously ^(34,35)^. Immunostaining was performed as described previously ^(36)^. The following primary antibodies were used including the products of Santa Cruz Biotechnology (Texas, USA): Ameloblastin, MMP20, DSP, K-Ras, FGF8, P21; ABCam: FGF4, FGF9, Caspase3, Ccnd1; Cell Signaling Tech (Massachusetts, USA): pSmad1/5/8, pP38, pErk1/2, pJNK, pAKT1, pPDK1; Merck Millipore: β-catenin; and Spring Bio: Ki67. Alexa Fluor 488 or 594 IgG (H+L) antibody (Thermo Fisher, Massachusetts, USA) and horseradish peroxidase-coupled IgG antibody (Santa Cruz) were used as the secondary antibody. For immunoblotting, pP38, pErk1/2, pJNK, pAkt, and Actin (Santa Cruz) were used as primary antibodies. IRDye 800cw or 680cw IgG (Odyssey, Nebraska, USA) were used as secondary antibodies. Immunoblotting was conducted as described previously ^(37)^. Signals were observed using microscope (Olympus BX51, Tokyo, Japan), Laser scanning confocal microscope (Zeiss LSM780, Jena, Germany), and Infrared Imaging Systems (Odyssey Clx). Representative images were taken from a minimum of 3 replicate experiments.

### Micro-computed tomography imaging and analysis

Harvested mineralized teeth were scanned with X-ray micro-computed tomography (microCT) equipment (μCT50, SCANCO Medical AG, Bruettisellen, Switzerland). The microCT parameters were as follows: 19 × 19 mm-field of view, 15-µm voxel size, 70 kVp, 200 mA, 360° rotation, 1500ms exposure time. All datasets were exported in the DICOM format with the isotropic 15-µm^3^ voxel. Acquired height and width of the tooth crown were analyzed with the attached software of μCT50.

### Statistics analysis

Image J software (version 1.46r; National Institutes of Health, USA) was utilized for Ki67-positive cell counting and to detect the densitometry of the western blot bands with at least three biological replicates counted. Statistical analyses were performed with GraphPad Prism 5 (Version 5.01; GraphPad Software Inc, California, USA). Data were represented as mean ± SD. Paired *t*-test was used to evaluate differences between two groups, and analysis of variance (ANOVA) was used to determine differences among different groups. If there was a significance of ANOVA test, Bonferroni post-tests was used. *p* values <0.05 is considered statistically significant difference.

## Results

### Tooth developmental pace is dominated by the dental mesenchyme in humans and mice

Our previous studies have shown that epithelial sheets of human keratinocyte stem cells, when recombined with E13.5 mouse dental mesenchyme, can be induced to differentiate into enamel-secreting ameloblasts in only 10 days and apoptotically degraded in 4 weeks, whereas the mouse secondary branchial arch epithelium, when recombined with the human dental mesenchyme from the bell-stage, remains in an enamel-secreting status after 15-week *ex vivo* culture ^(30,38,39)^. These results strongly imply that the duration of epithelial differentiation in tooth development could be dominated by the mesenchymal component in humans and mice. To validate our hypothesis, we comprehensively revisited these phenotypes by carrying out meticulously designed heterogenous recombination between human and mouse embryonic dental tissues (Table 1). As shown in Fig. 1A-C, it took more than 4 weeks for the mouse dental epithelium to deposit enamel when recombined with the human dental mesenchyme in mde+hdm chimeric tooth germs and the chimera generated a mineralized tooth after 16-week growth (Suppl. Fig. 1A), exhibiting a much longer developmental duration and slower tempo in comparison with the mouse dental epithelia that were combined with their own dental mesenchyme (Fig. 1D-F, Suppl. Fig. 1B). By contrast, it took only about 10 days for the human dental epithelium to deposit enamel when recombined with E13.5 mouse dental mesenchyme in hde+mdm chimeric tooth germs (Fig. 1G-I) and the chimera could achieve a mineralized tooth in less than 30 days as well as the mouse homogenous (mde+mdm) tooth (Suppl. Fig. 1C), demonstrating a dramatically accelerated developmental pace compared to the normal human tooth morphogenesis. Immunostaining of two ameloblastic differentiation markers, ameloblastin and MMP20 (Matrix Metallopeptidase 20), further confirmed the development phase alteration of the dental epithelium induced by the heterogenous dental mesenchyme. The expression of these two markers in the mouse dental epithelium-derived pre-ameloblasts in mde+hdm chimeric recombinants was delayed to 4^th^ week in subrenal culture (Fig. 1J-L), whereas the expression of the two markers was antedated at 9-day in the human dental epithelium-derived ameloblasts in hde+mdm chimeric recombinants in subrenal culture (Fig. 1P-R), which is similar to the expression of them at 7-day in the ameloblasts derived from the mouse dental epithelium in mde+mdm control recombinants (Fig. 1M-O). The possibility of tissue contamination was ruled out using specific primary antibodies against mouse or human MHC I antigens to distinguish between human and mouse tissues in the heterogenous chimeras (Suppl. Fig. 2). Our results unambiguously indicate that the dental mesenchyme is a crucial component that dominates the tooth developmental pace by interaction with the dental epithelium via diffusible paracrine signals during odontogenesis.

**Fig. 1.**
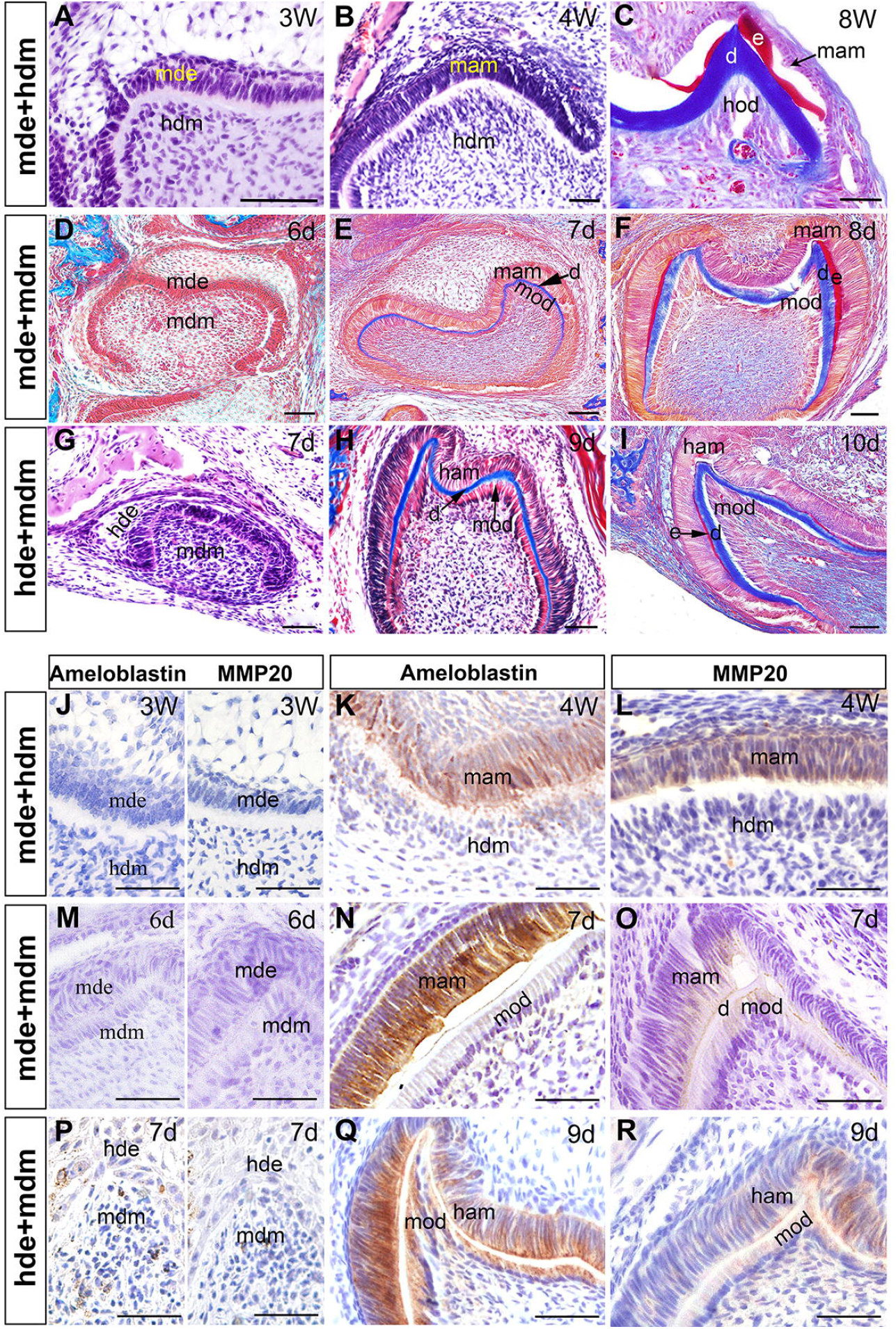
Heterogenous recombination of human and mouse embryonic dental tissues reveals the dominant role of the dental mesenchyme in the control of tooth developmental pace. **A-C**: Azan dichromic staining (stains dentin blue and enamel red) of recombinants of E13.5 mouse dental epithelium and bell-stage human dental mesenchyme cultured under the kidney capsule for 3 weeks (**A**), 4 weeks (**B**), 8 weeks (**C**). Note that no deposition of dentin and enamel was seen in the chimeric tooth germs until 4 weeks in subrenal culture. **D**-**F**: Azan dichromic staining of recombinants of E13.5 mouse dental epithelium and E13.5 mouse dental mesenchyme cultured under the kidney capsule for 6 days (**D**), 7 days (**E**), and 8 days (**F**). Note that the deposition of dentin and enamel was not seen until 7 and 8 days, respectively, in subrenal culture. **G**-**I**: Azan dichromic staining of recombinants of bell-stage human dental epithelium and E13.5 mouse dental mesenchyme cultured under the kidney capsule for 7 days (**G**), 9 days (**H**), and 10 days (**I**). Note that the deposition of dentin and enamel was not seen until 9 and 10 days, respectively, in subrenal culture. **J-R**: The expression of ameloblastin and MMP20 in recombinants of E13.5 mouse dental epithelium and bell-stage human dental mesenchyme cultured under the kidney capsule for 3 weeks (**J**), 4 weeks (**K** and **L**), in recombinants of E13.5 mouse dental epithelium and mouse dental mesenchyme for 6 days (**M**), 7 days (**N** and **O**), and in recombinants of bell-stage human dental epithelium and mouse dental mesenchyme for 7 days (**P**) and 9 days (**Q** and **R**). hdm, human dental mesenchyme; hde, human dental epithelium; mdm, mouse dental mesenchyme; mde, mouse dental epithelium; hod, human odontoblasts; mod, mouse odontoblasts; mam, mouse ameloblasts; d, dentin; e, enamel. Scale bar = 50 μm.

### FGF signaling participates in the regulation of tooth developmental pace

We next attempted to identify the paracrine signaling molecules that may be involved in the regulation of tooth developmental tempo in humans and mice. Conserved SHH, FGF, BMP, and Wnt pathways are all crucial signaling engaged in cell proliferation, apoptosis and differentiation during odontogenesis, indicating that they may be key players in orchestrating the fundamental mainspring of organ growth. Since the dental mesenchyme is dominant in the regulation of tooth growth (Fig. 1), we exclude epithelially expressed SHH ^(40)^. To test whether mesenchymal BMP, FGF and WNT signaling may be involved in the regulation of tooth developmental pace, we examined the level of the active indicators of these pathway including pSmad1/5/8 for BMP, K-Ras for FGF, and β-catenin for Wnt, respectively, in the chimeric tooth germs. The expression level of K-Ras in both the dental epithelium and mesenchyme is much higher in the mde+hdm chimeric tooth germ (Fig. 2E), where the development of mouse dental epithelium-derived ameloblasts were dramatically delayed and remained immature after 4 weeks in subrenal culture, compared to that in the mouse homogenous recombinant tooth germ (Fig. 2B). The expression levels of pSmad1/5/8 and β-catenin in mouse dental epithelium-derived ameloblasts and adjacently interacted odontoblasts did not display obvious difference as compared between the heterogenous recombinants and homogenous ones (Fig. 2A,C,D,F). The enhanced epithelial FGF activity in the mde+hdm chimeric tooth germ, evidenced by outstandingly enhanced activity of K-Ras, was certainly induced by the human dental mesenchyme and may attribute to the protracted differentiation of the mouse dental epithelium.

**Fig. 2.**
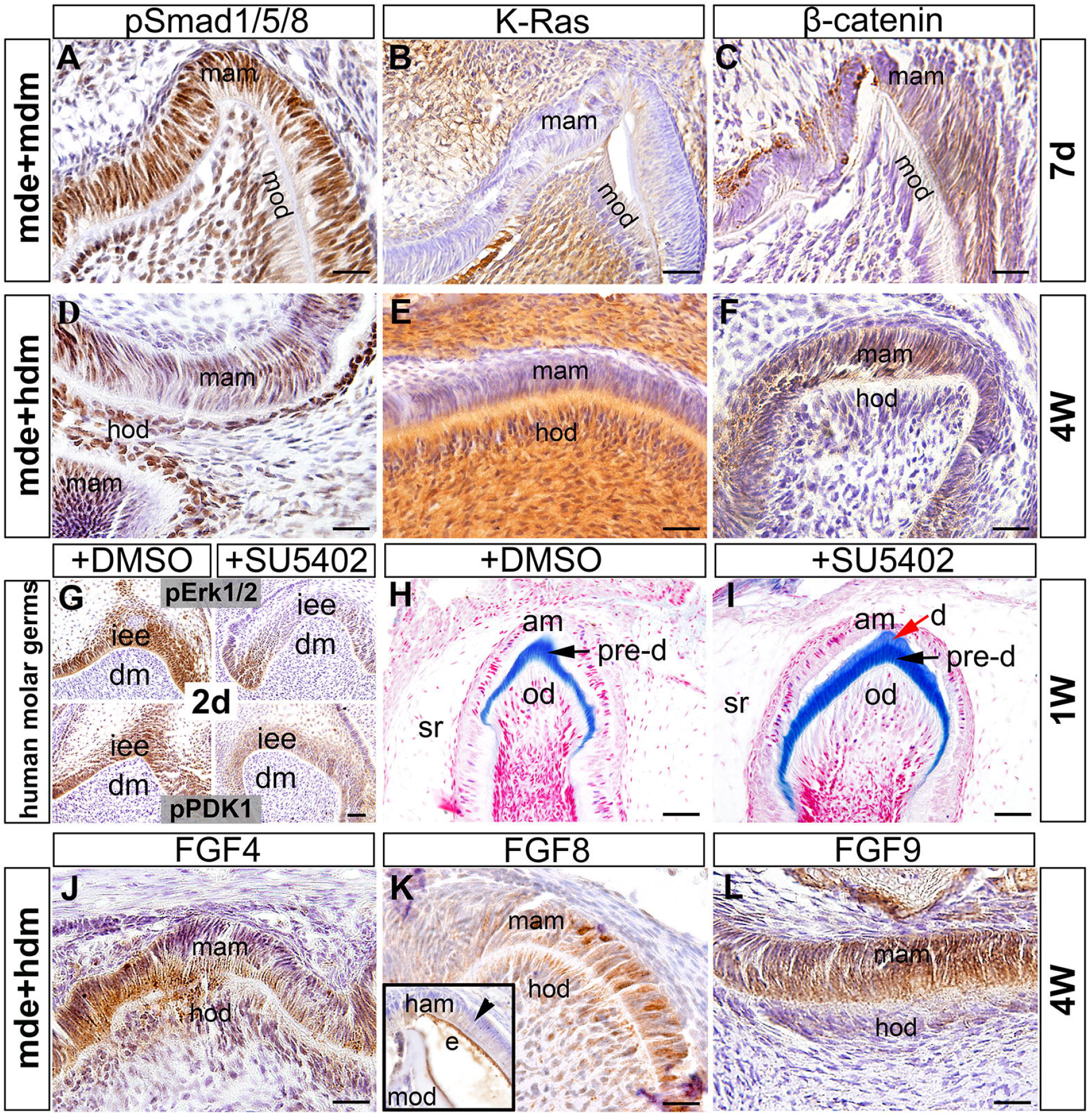
FGF signaling is involved in the regulation of tooth developmental pace. **A-C**: Immunostaining showing the expression of pSmad1/5/8 (**A**), K-Ras (**B**), and β-catenin (**C**) in recombinants of mouse dental epithelium and mouse dental mesenchyme cultured under the kidney capsule for 7 days. **D-F**: Immunostaining showing the expression of pSmad1/5/8 (**D**), K-Ras (**E**), and β-catenin (**F**) in recombinants of mouse dental epithelium and human dental mesenchyme cultured under the kidney capsule for 4 weeks. **G**: Human 1^st^ premolar germs of 16^th^ week gestation were transplanted under the kidney capsule and cultured in the absence or presence of SU5402 for 2 days. Note the decreased expression of pErk1/2 and pPDK1 in the presence of SU5402. **H**-**I**: Human 1^st^ premolar germs of 16^th^ week gestation were transplanted under the kidney capsule and cultured in the absence (**H**) or presence (**I**) of SU5402 for 1 week. Note the accelerated detinogenesis with the appearance of dentin (**I**, red arrow) in the presence of SU5402. **J-L**: Immunohistochemical staining showing the expression of FGF4 (**J**), FGF8 (**K**), and FGF9 (**L**) in recombinants of mouse dental epithelium and human dental mesenchyme cultured under the kidney capsule for 4 weeks and the expression of FGF8 (insert in **K**) in recombinants of human dental epithelium and mouse dental mesenchyme cultured under the kidney capsule for 10 days. mam, mouse ameloblasts; mod, mouse odontoblasts; hod, human odontoblasts; am, ameloblasts; od, odontoblasts; iee, inner enamel epithelium; sr, stellate reticulum; dm, dental mesenchyme; d, dentin; pre-d, predentin. Scale bar = 25 μm.

To explore whether the upregulated FGF signaling may be associated with the prolonged tooth development phase in humans, we started with a search for candidate FGFs that are persistently expressed in the human developing dental mesenchyme but barely in the mouse one. Most expression patterns of 22 known FGF ligands in mammals have been verified in mouse tooth germs at the cap and bell stages (summarized in Suppl. Table 1)^(41,42)^. We have examined in our previous studies the expression patterns of *FGF3, FGF4, FGF7, FGF8, FGF9*, and *FGF10* in human tooth germs at different stages using *in situ* hybridization or immunohistochemistry ^(43)^. Here, we screened the expression of additional 12 *FGFs* that have not been identified in human tooth germs by RT-PCR. We excluded *FGF11, FGF12, FGF13*, and *FGF14*, because they are not secreting ligands and do not activate FGF receptors as well. We found that, among them, the transcripts of *FGF1, FGF2, FGF15/19*, and *FGF18* were detectable at both the cap and bell stages (data not shown). We further confirmed their expression in developing human teeth by section *in situ* hybridization (Suppl. Fig. 3). When comparing the FGF expression data between the human ^(43)^ and mouse ^(41,42)^, we noticed that the majority of FGF transcripts are broadly distributed in the whole tooth germ including the dental epithelium and mesenchyme throughout the early cap stage to the late bell stage in humans ^(43)^. However, their expression is restricted to the limited region of tooth germs, such as enamel knot, and appeared in limited periods of time in mice (Suppl. Table 1)^(41,42)^. We believe that this widely and persistently distributing pattern of FGF ligands in the human tooth germs is closely related to their characteristics of tooth development. To further verify this notion, we cultured human 1^st^ premolar germs at the early bell stage under the kidney capsule of nude mice with intraperitoneal injection of SU5402, a global FGF-signaling inhibitor (an inhibitor of tyrosine phosphorylation of FGF receptor). The inhibition activity of SU5402 in human tooth germs was firstly evaluated after two days injection and showed that it significantly inhibited FGF-mediated activation of FGFR downstream signal transduction, pErk1/2 and pPDK1 (Fig. 2G). Then, we cultured the human molar germs with SU5402 for one week and found accelerated detinogenesis in the SU5402-treated molar germ with the appearance of both predentin and dentin (Fig. 2I), as compared to DMSO-treated one with appearance of only predentin (Fig. 2H), indicating pivotal roles of FGF-signaling in regulation of tooth developmental tempo.

With further analyses of FGF expression data in humans and mice, we found that FGF4, FGF8, and FGF9 ligands exhibit most distinctive patterns (Suppl. Table 1). These three ligands are persistently expressed in the dental epithelium and mesenchyme at least until the differentiation stage in humans ^(43)^. However, in mice, the expression of FGF4 is restricted to the enamel knot from the cap to bell stage and that of FGF9 to the enamel knot at the cap stage and to the dental epithelium at the bell stage. Furthermore, FGF8 is only expressed in the presumptive dental epithelium prior to and during the laminar stage and is no longer expressed after the E12 bud stage ^(44)^. Whereas, in humans, *FGF8* transcripts are widely and intensively present in the whole tooth germ at least from the early cap stage to the differentiation stage ^(43)^. Importantly, we found retained expression of FGF8, as well as FGF4 and FGF9, in the mouse dental epithelium derived pre-ameloblasts in mde+hdm recombinants after 4 weeks in subrenal culture, where the development of mouse ameloblasts was largely delayed (Fig. 2J-L), whereas FGF8 was attenuated in the human dental epithelium derived pre-ameloblasts in hde+mdm recombinants after 10 days in subrenal culture, where the development of human ameloblasts was vastly accelerated (insert in Fig. 2K). Based on these observations, we hypothesized that FGF signaling may play important roles and FGF8 could be a critical factor in the regulation of tooth developmental pace.

### Forced activation of FGF8 signaling in the mouse dental mesenchyme delays tooth development

To test our hypothesis, we set to overexpress *Fgf8* in dental mesenchymal cells in the E13.5 recombinant molar germ via lentivirus-mediated expression and subrenal culture system. Histological examination revealed that it took up to 9 days for *Fgf8*-overexpressed tooth grafts to start dentin secreting (Fig. 3B), whereas it only took 7 days for control ones to do so (Fig. 3A). Moreover, Ki67 staining detected remarkably higher level of proliferation rate in both the dental epithelium and mesenchyme of *Fgf8*-overexpressed tooth grafts compared to the controls after 7 days in subrenal culture (Fig. 3C-E; *n*=6/group). These data indicate that activated mesenchymal FGF8 activity delays tooth development and dramatically stimulates cell proliferation.

**Fig. 3.**
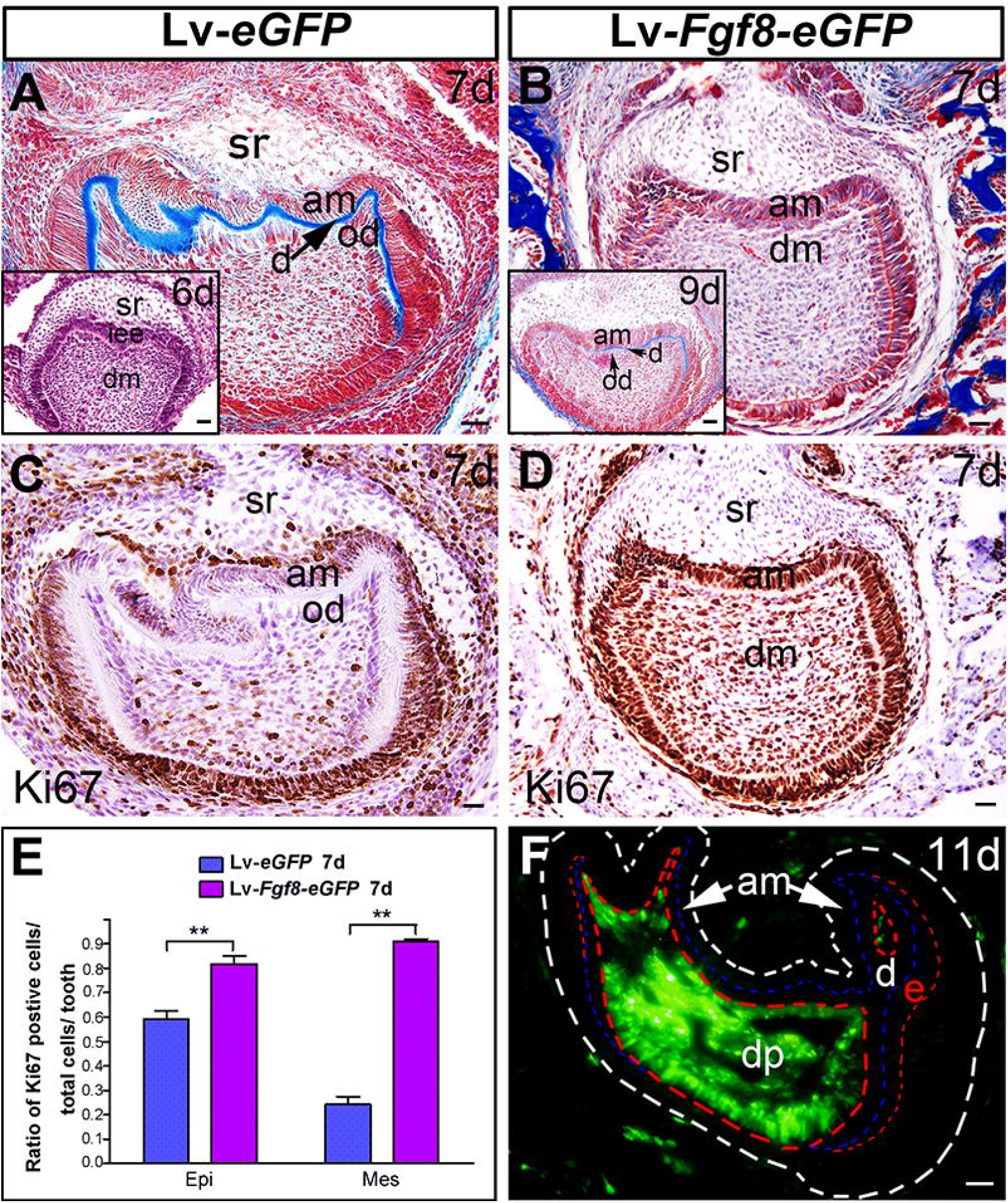
Augmented FGF8 signaling in the mouse dental mesenchyme delays tooth development and increases cell proliferation. **A, B**: Azan dichromic staining of recombinants of E13.5 mouse dental epithelium and re-aggregate of dental mesenchymal cells infected with control LV*-eGFP* (**A**) or LV-*Fgf8-eGFP* (**B**) cultured under the kidney capsule for 6 days (insert in **A**), 7 days, and 9 days (insert in **B**). **C, D**: Immunostaining showing the expression of Ki67 in recombinants of E13.5 mouse dental epithelium and re-aggregate of dental mesenchymal cells infected with control LV*-eGFP* (**C**) or LV-*Fgf8-eGFP* (**D**) cultured under the kidney capsule for 7 days. **E**: Comparison of Ki67-positive cells in the dental mesenchyme and dental epithelium of the recombinants. Standard deviation values were shown as error bars, and ** indicates *p*<0.01, *n* = 6/group. **F**: GFP staining of recombinants of E13.5 mouse dental epithelium and re-aggregate of dental mesenchymal cells infected with control LV*-eGFP* cultured under the kidney capsule for 11 days. dm, dental mesenchyme; dp, dental pulp; am, ameloblasts; sr, stellate reticulum; od, odontoblasts; d, dentin; Epi, epithelium; Mes, mesenchyme. Scale bar = 25 μm.

To corroborate the aforementioned *ex vivo* results, we further generated *Wnt1-Cre; R26R*^*Fgf8*^ transgenic mice to verify whether forced activation of mesenchymal FGF8 signaling alters tooth development *in vivo*. The presence of FGF8 in the dental mesenchyme of *Wnt1-Cre;R26R*^*Fgf8*^ mice was confirmed by anti-FGF8 immunostaining (Suppl. Fig. 4). Most of mutant embryos die at around E12.5 and a few died around E14.5. Although histological examination revealed severe craniofacial hypoplasia in 100% cases, including exencephaly and the absence of identifiable tissues such as ears, auricles, eyes, nose, as well as tongue (data not shown), the mutant mice still possessed a pair of intact maxillary and mandibular molar germs (Fig. 4). More importantly, both the maxillary and mandibular molar germ exhibited an obvious retardation of tooth development (Fig. 4). Since most mutant embryos died at around E12.5, we grafted E11.5 mutant molar germs under the kidney capsule for further development. Examination of histology and the deposition of dentin and enamel showed that the mutant maxillary molar developed with approximate one-day’s delay when compared to the wild-type ones in nearly 100% (15/16, Fig. 5A-F), and the mutant mandibular molar developed with about two-day’s delay in 2 out of 9 grafts with the rest 7 failing to develop (insert in Fig. 5D). Examination of the expression of odontoblast marker DSP and ameloblast marker ameloblastin in the grafted maxillary molars further confirmed this delay (Fig. 5I-L).

**Fig. 4.**
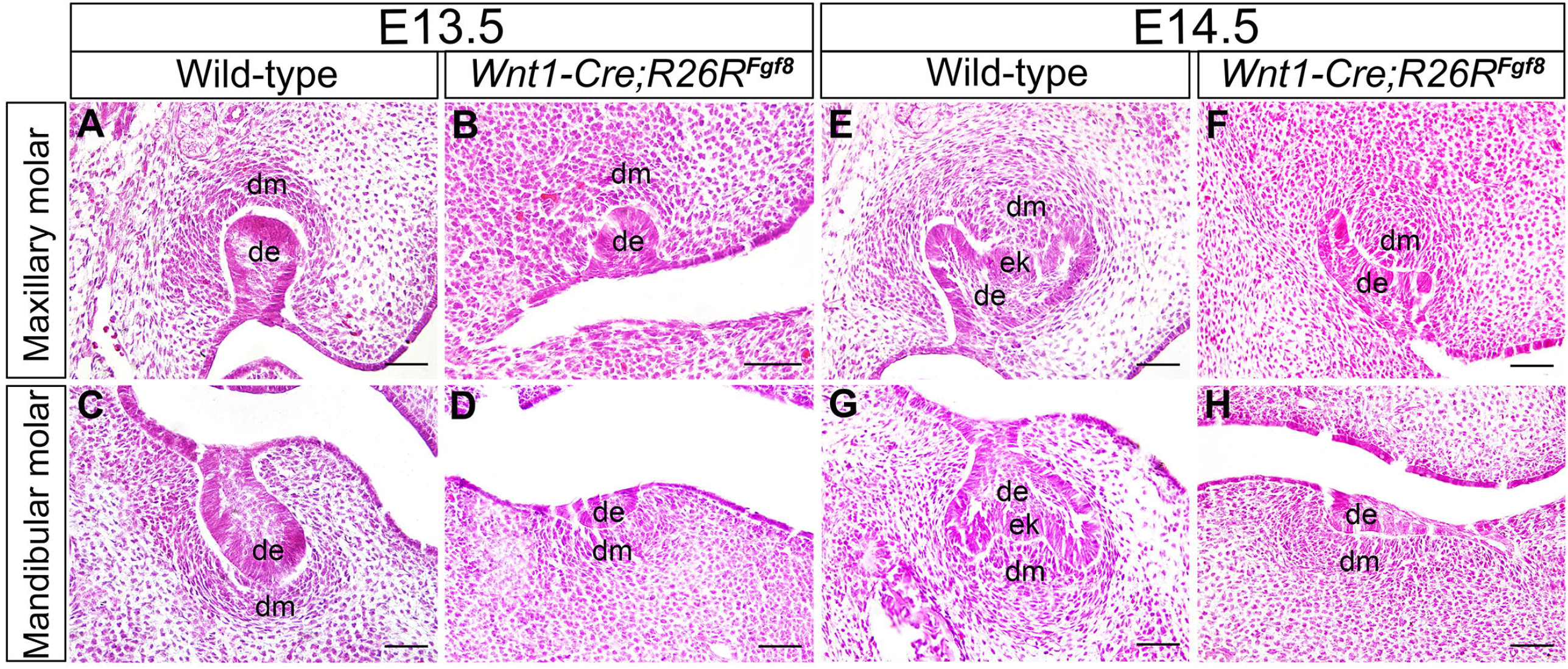
Delayed tooth development in early molar germs of *Wnt1-Cre;R26R*^*Fgf8*^ mice. **A-H:** H&E staining showing the morphology of maxillary and mandibular molar in E13.5 wild-type (**A**,**C**) and mutant (**B**,**D**) embryos and in E14.5 wild-type (**E**,**G**) and mutant (**F**,**H**) embryos. de, dental epithelium; dm, dental mesenchyme. Scale bar = 50 μm.

**Fig. 5.**
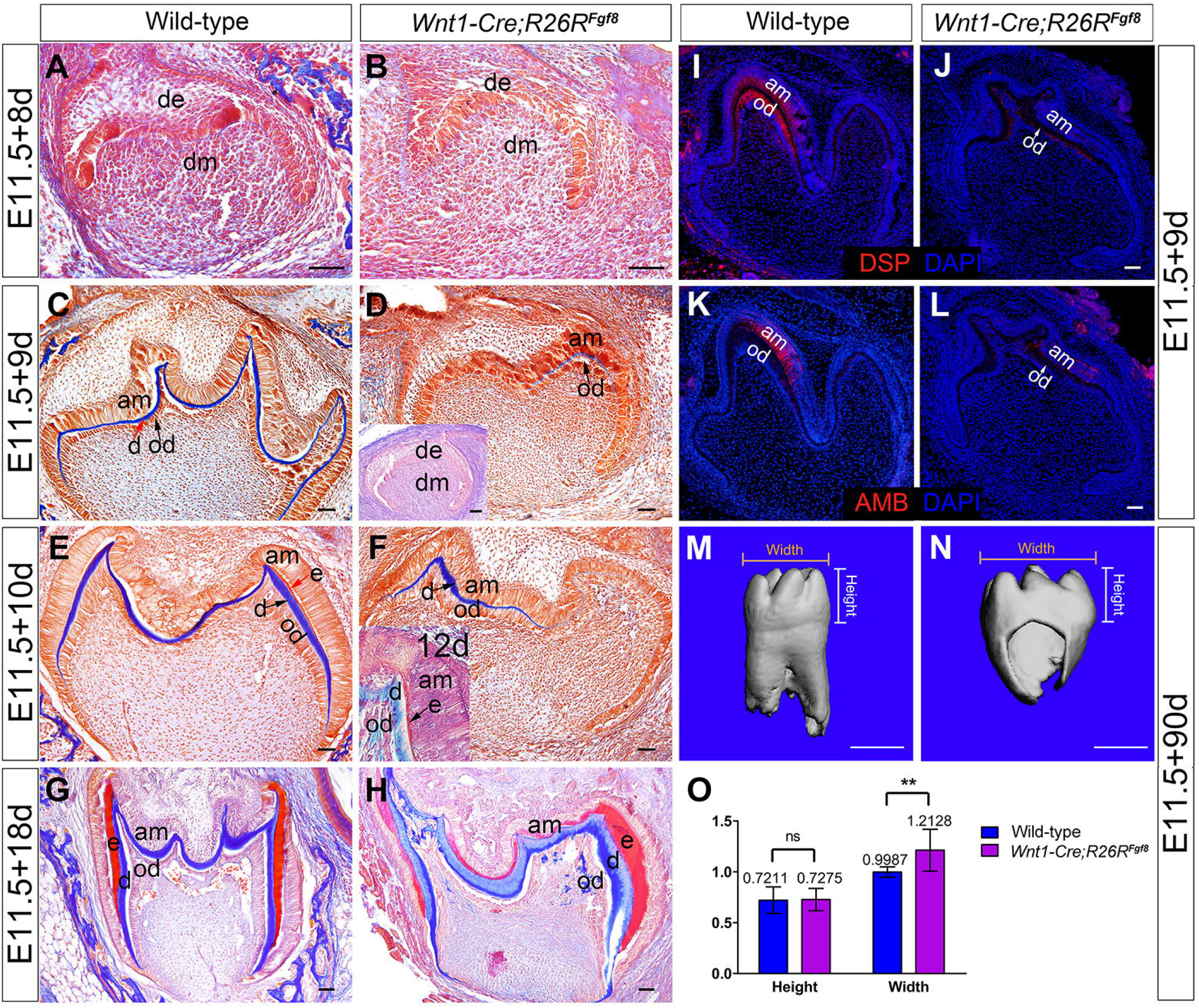
Grafts of *Wnt1-Cre;R26R*^*Fgf8*^ molar germs exhibit delayed tooth development and enlarged tooth size. **A-H**: Azan dichromic staining of E11.5 wild-type and mutant maxillary molar germs cultured under the kidney capsule for 8 days (**A**,**B**), 9 days (**C**,**D**), 10 days (**E**,**F**), 12 days (insert in **F**), and 18 days (**G**,**H**). Azan dichromic staining of E11.5 mutant mandibular molar germs cultured under the kidney capsule for 9 days (insert in **D**). **I-L**: Immunostaining showing the expression of DSP (**I**,**J**) and Ameloblastin (**K**,**L**) in E11.5 wild-type and mutant molar grafts cultured under the kidney capsule for 9 days. **M, N**: Representative Micro-CT images of E11.5 wild-type (**M**) and mutant molar germs (**N**) cultured under the kidney capsule for 3 months. **O**: Comparison of the height and width of the tooth crown between wild-type and mutant mineralized teeth cultured under the kidney capsule for 3 months. Standard deviation values were shown as error bars. Ns indicates *p* >0.05, and ** indicates *p*<0.01, *n* = 7 and 11/group in **O**. de, dental epithelium; dm, dental mesenchyme; am, ameloblasts; od, odontoblasts; d, dentin; e, enamel. Scale bar: A-H, I-L = 50 μm; M and N = 500 μm.

Interestingly, Azan staining showed that the mutant graft of maxillary molar germ formed a larger-sized tooth (Fig. 5G) than the wild-type one (Fig. 5H) after 18 days in subrenal culture. Further Micro-CT scanning revealed that the mutant graft indeed formed a larger-sized tooth with short and dumpy shape (Fig 5N) compared to the control (Fig. 5M) after 3 months in subrenal culture for complete mineralization, which exhibited no difference in the average height of the tooth crowns but about 20% wider in the average width in the mutants (*n* = 11) compared to the controls (*n* = 7, Fig. 5O). Apparently, higher rate of cell proliferation contributes greatly to the larger teeth, which was evidenced by more Ki67 positive cells presented in E13.5 mutant tooth germ and the grafted mutant teeth than controls (Fig. 6A-F, K-M, *n* = 3/each group). Meanwhile, while apoptotic marker Caspase3 staining was present in ameloblasts and odontoblasts in both mutant (Fig. 6I,J) and wild-type tooth germs (Fig. 6G,H), this apoptotic signal was non-detectable in the pulp cells of mutant tooth germs (Fig. 6I,J), whereas it was seen in that of wild-type ones (Fig. 6G,H) after 18 days in subrenal culture. These results further support the idea that ectopically activated FGF8 signaling in the mouse dental mesenchyme stimulates cell proliferation and prevents cell apoptosis during odontogenesis.

**Fig. 6.**
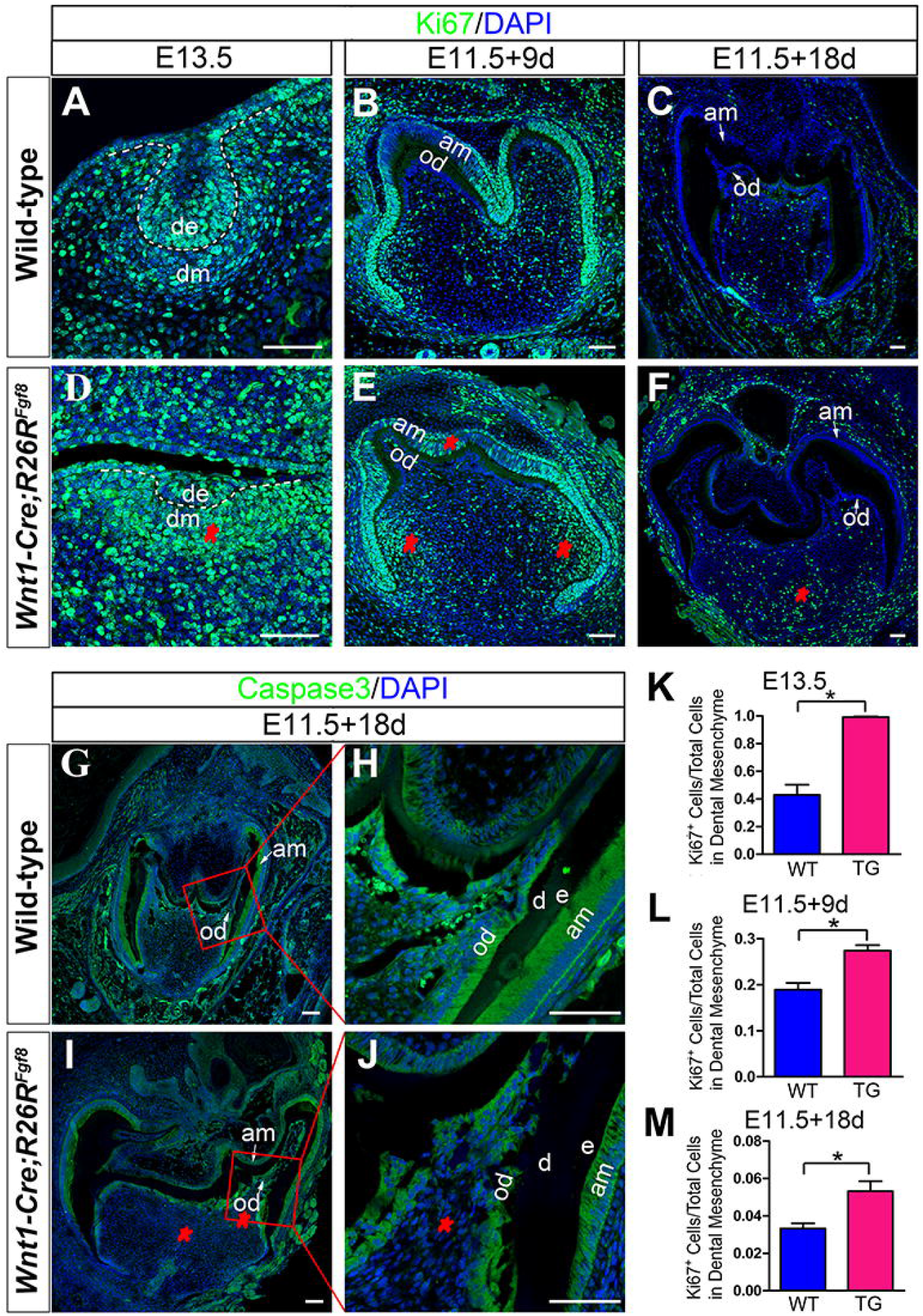
Ectopically activated dental mesenchymal FGF8 signaling stimulates cell proliferation and prevents cell apoptosis during odontogenesis. **A**-**F**: Immunostaining showing the Ki67 positive cells in E13.5 wild-type (**A**) and mutant (**D**) molar germs and in the grafts of E11.5 wild-type and mutant molar germ cultured under the kidney capsule for 9 days (**B**,**E**) and 18 days (**C**,**F**). **G-J**: Immunostaining showing the expression of Caspase3 in the grafts of E11.5 wild-type (**G**, boxed area is magnified in **H**) and mutant (**I**, boxed area is magnified in **J**) molar germ cultured under the kidney capsule for 18 days. Note the absence of Caspase3 expression in the dental mesenchyme (red asterisk) excepting odontoblasts in mutant grafts. **K-M**: The percentage of Ki67 positive cells in the dental mesenchyme of wild-type and mutant E13.5 tooth germs (**K**) and tooth grafts cultured under the kidney capsule for 9 days (**I**) and 18 days (**M**), *n* = 3/group, * indicates *p* < 0.05. Standard deviation values were shown as error bars. am, ameloblasts; od, odontoblasts; d, dentin; e, enamel. Scale bar = 100 μm.

### Ectopically activated dental mesenchymal FGF8 signaling is transduced via p38 and PI3K-Akt intracellular pathway

To determine which intracellular pathway is implicated in the signal transduction of the ectopically activated dental mesenchymal FGF8 signals, we examined the levels of phosphorylated Erk1/2, P38, JNK, and PI3K-Akt, the activity indicator of MAPK pathway, in both E13.5 tooth germs and E11.5 ones after 9 days in subrenal culture. As shown in Fig. 7, no signal intensity differences in pErk1/2 (Fig. 7A-D) and pJNK (Fig. 7E-H) staining were observed between wild-type and mutant teeth. Whereas, p38 signaling is dramatically elevated in *Wnt1-Cre;R26R*^*Fgf8*^ teeth with a considerably increased amount of nuclear-localized pP38 cells both in the dental epithelium and dental mesenchyme of E13.5 mutant tooth germs (Fig. 7J) and grafted teeth (Fig. 7L) as compared to the controls (Fig. 7I,K). Meanwhile, the intensity of PI3K-Akt staining was slightly enhanced and the amount of pAkt1^+^ cells was increased in mutant E13.5 tooth germs and grafted teeth (Fig. 7N,P) as compared to controls (Fig7 M,O). Western blot analysis further confirmed the increased expression of pP38 and pAkt1 in E13.5 mutant tooth germs (Fig. 7Q,R; *n* = 3). Our results indicate that the augmented dental mesenchymal FGF8 activity enhances P38 and PI3K-Akt signaling in both the dental mesenchyme and the dental epithelium, suggesting that they may function as intracellular pathways for transducing FGF8 signal into nuclei.

**Fig. 7.**
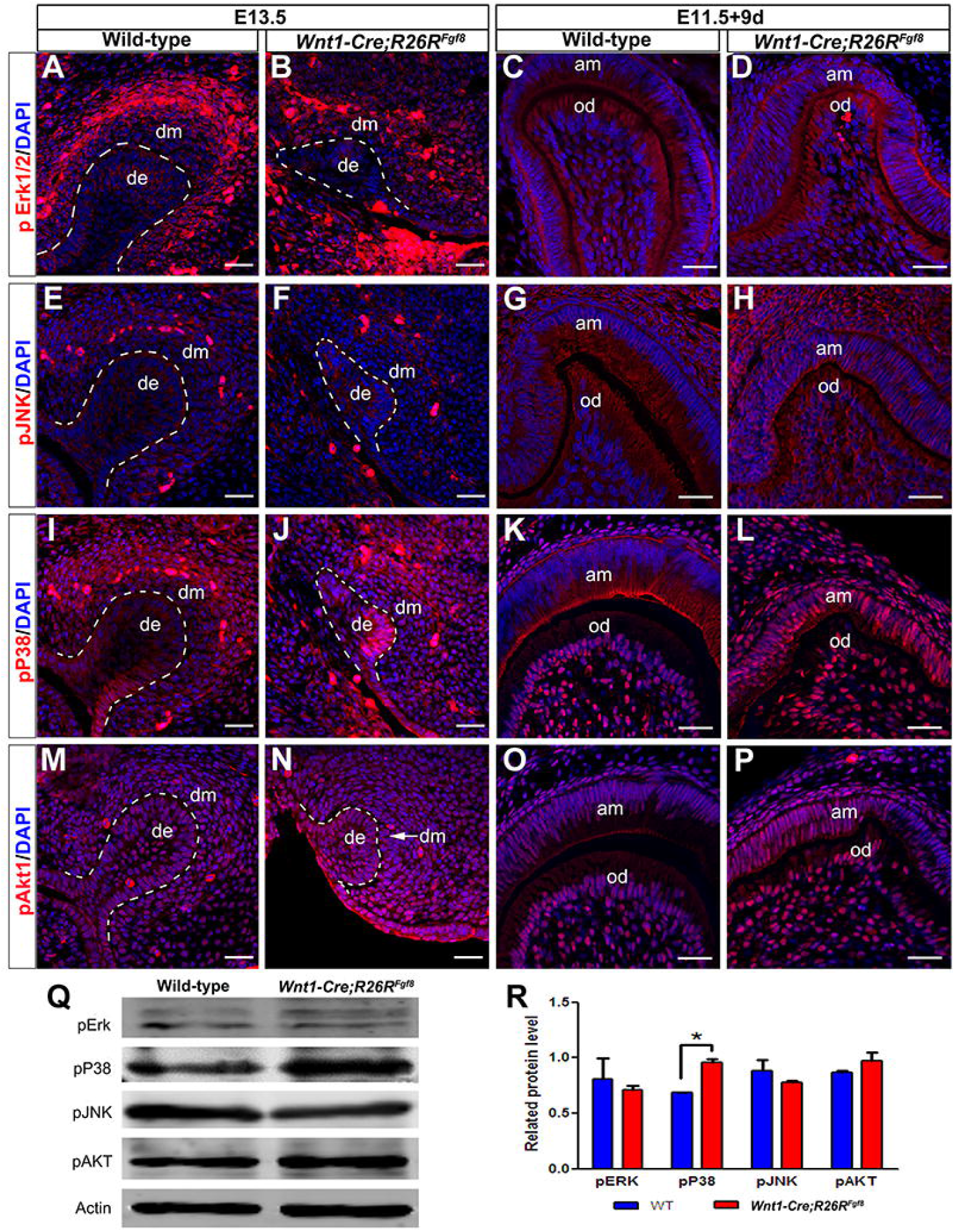
Ectopically activated dental mesenchymal FGF8 signaling is transduced via p38 and PI3K-Akt intracellular pathways. **A-D**: Immunostaining showing the expression of pErk1/2 in E13.5 wild-type (**A**) and mutant (**B**) molar germs and in E11.5 wild-type (**C**) and mutant (**D**) molar germs cultured under the kidney capsule for 9 days. **E-H**: Immunostaining showing the expression of pJNK in E13.5 wild-types (**E**) and mutants (**F**) molar germs and in E11.5 wild-type (**G**) and mutant (**H**) molar germs cultured under the kidney capsule for 9 days. **I-L**: Immunostaining showing the expression of pP38 in E13.5 wild-type (**I**) and mutant (**J**) molar germs and in E11.5 wild-type (**K**) and mutant (**L**) cultured under the kidney capsule for 9 days. **M-P**: Immunostaining showing the expression of pAkt1 in E13.5 wild-types (**M**) and mutants (**N**) molar germs and in E11.5 wild-type (**O**) and mutant (**P**) molar germs cultured under the kidney capsule for 9 days. **Q**: Western-blotting of pErk1/2, pP38, pJNK, and pAKT1 expressed in E13.5 wild-type and mutant molar germs. **R**: Quantification of immunoblotting data in **Q**. *n* = 3/group. * indicates *p <* 0.05. Standard deviation values were shown as error bars. de, dental epithelium; dm, dental mesenchyme; am, ameloblasts; od, odontoblasts. Scale bar = 25 μm.

### Enhanced dental mesenchymal FGF8 signaling promotes transition of cell cycle from G1-to S-stage during odontogenesis

The rate of cell proliferation controlled by cell cycle progression is a key player in organ size control ^(4)^ and is also closely associated with tissue differentiation status, as higher cell proliferation always presents in developing tissues and lower one in differentiated ones. Since the elevation of *Fgf8* expression in the dental mesenchyme results in the significant promotion of cell proliferation and prevention of cell apoptosis (Fig. 6), we speculated that enhanced dental mesenchymal FGF8 signaling would alter cell cycling. To verify this hypothesis, we set forth to ascertain the distribution of cyclin D1 (Ccnd1) and its inhibitor p21^Cip1^ (p21), a pair of critical regulators functioning stimulatory and repression effect, respectively, on inducing cell cycle entry and mediating the G1-S stage transition ^(45,46)^, in E13.5 tooth germs and grafted teeth. As shown in Fig. 8, although the same high density of Ccnd1^+^ cells were present in the dental epithelium of E13.5 wild-type and mutant tooth germs, the dental mesenchyme in mutant tooth germs possessed a considerably increased amount of Ccnd1^+^ cells in comparison to that in wild-type ones (Fig. 8A,E). On the other hand, p21^+^ cells were abundantly present in the presumptive primary enamel knot of the dental epithelium and slightly in the dental mesenchyme in wild-type tooth germs but totally absent from the whole tooth germ in mutant ones (Fig. 8B,F). Thereout, more colocalization of Ccnd1 and p21 staining were found in the wild-type tooth germs than in the mutant ones (Fig. 8C-D, G-H). In grafted teeth, nuclear Ccnd1^+^ cells were distributed not only in ameloblasts and odontoblasts (Fig. 8M,P-upper) but also all over the dental pulp in mutants (Fig. 8M,P-lower), rather than being restricted to the dental mesenchymal cells adjacent to the odontoblasts in wild-types (Fig. 8I,L). Although p21 was expressed in ameloblasts and dental mesenchymal cells in both wild-type and mutant teeth, its expression level was relatively decreased in mutants (Fig. 8J,L,N,P). In addition, the nuclear localization of p21 could be detected in dental mesenchymal cells of wild-type teeth (Fig. 8L-lower), whereas it was found to be in the cytoplasm of grafted mutant teeth (Fig. 8P-lower), which make it unable to cooperate with nuclear Ccnd1 to regulate the cell cycle progression in the dental mesenchymal cells. Taken together, these results provide evidence that enhanced mesenchymal FGF8 activity observably elevates the activity of nuclear Ccnd1 while degrades the activity of nuclear p21. Thus both stimulation of Ccnd1 and repression of p21 function to induce cell cycle entry and promote the transition of cell cycle from G1-to S-stage, which facilitates cell population to retain the high rate of proliferation.

**Fig. 8.**
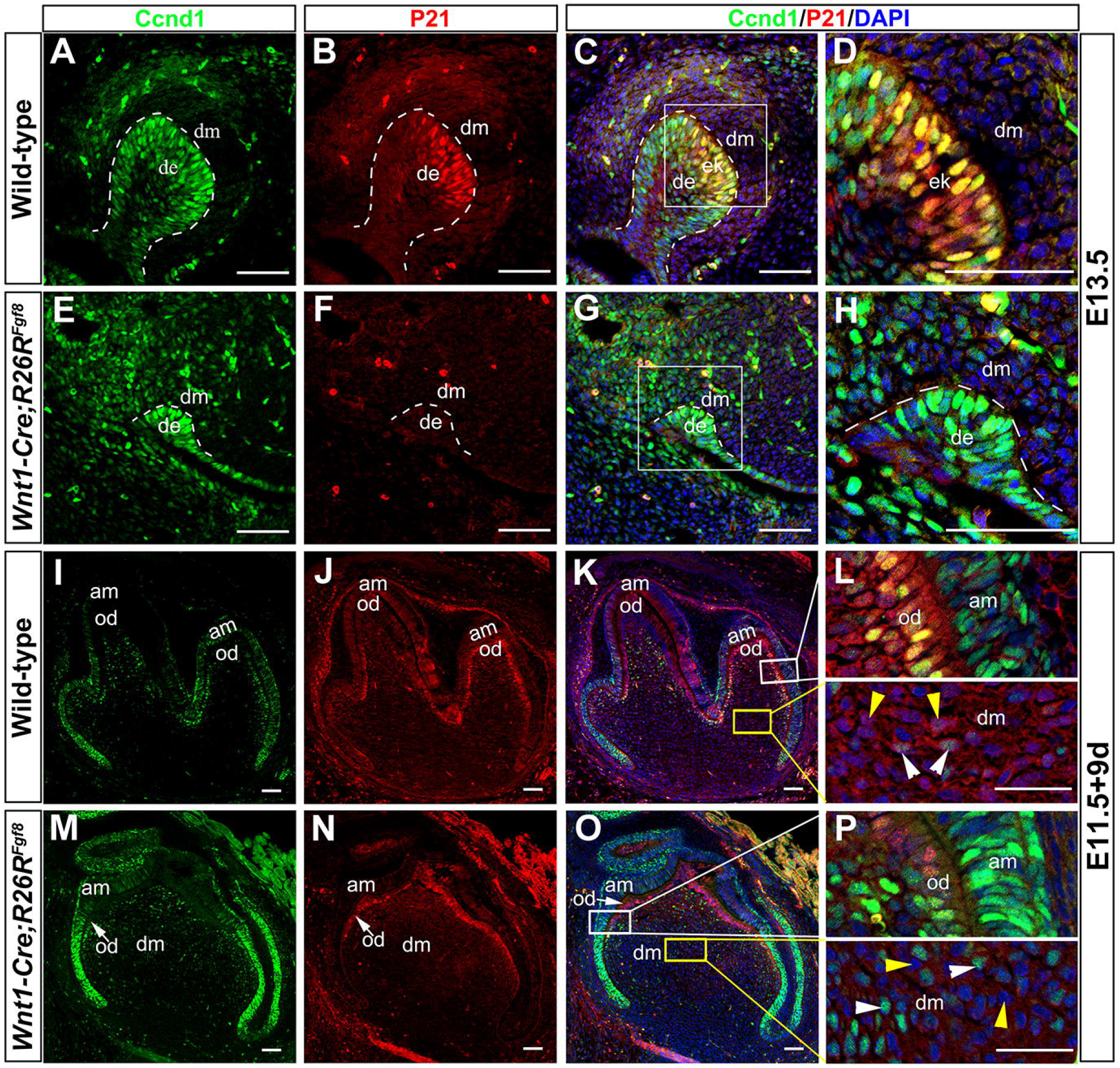
Enhanced dental mesenchymal FGF8 signaling increases the level of nuclear Ccnd1 and decreases the level of nuclear p21. Immunostaining showing the colocalization of Ccnd1 and P21 in E13.5 wild-type (**A-D**) and mutant (**E-H**) molar germs and E11.5 wild-type (**I-L**) and mutant (**M-P**) molar germs cultured under the kidney capsule for 9 days. Boxed areas in (**C**), (**G**), (**K**), and (**O**) are magnified in (**D**), (**H**), (**L**), and (**P**), respectively. Note the significantly increased proportion of Ccnd1-positive cells (white arrow) and the shift of P21 from the nuclei to the cytoplasm (yellow arrow) in the mutant dental mesenchyme (**P**) as compared with wild-type ones (**L**). de, dental epithelium; dm, dental mesenchyme; am, ameloblasts; od, odontoblasts. Scale bar: A-N, P-S = 50 μm; O, T = 25 μm.

## Discussion

### Tooth developmental tempo complies with the hereditary nature of dental mesenchyme

A heterogenous recombination study carried out between the mouse and rat molar tooth germs has suggested that tooth size is determined by the dental mesenchyme ^(47,48)^. Additionally, our recent studies showed that, in comparison with natural developmental pace, the differentiation duration of the mouse dental epithelium or non-dental epithelium into ameloblasts when induced by the human dental mesenchyme is significantly extended, whereas that of the human dental epithelium or epithelial stem cells into ameloblasts when induced by the mouse dental mesenchyme is significantly shortened ^(38,39)^. These results are consistent with the difference of development progression speed between human and mouse teeth ^(13)^, suggesting a dominant role of the dental mesenchyme in regulation of tooth developmental tempo. This idea is further supported by our current heterogenous recombination study carried out between the mouse and human molar tooth germs (Fig. 1). We further provide unambiguous evidence that not only the tooth size but also the tooth developmental pace are obedient to the hereditary character of dental mesenchyme.

### Dental mesenchymal FGF8 and synergistic effects of FGFs play critical roles in controlling the developmental tempo during mammalian odontogenesis

Conserved pathways, such as FGF, BMP, Hh, and WNT pathway, participate in the highly conserved molecular mechanism underlying organismal growth by corresponding alterations of cell growth and division ^(4,11,12)^. FGFs, as the most common mitogens, are positively correlated to the rate of cell growth and cell division and involved in the induction and pattern formation during tooth development ^(49,50)^. Fgf8 has been demonstrated to play an important role in cell survival and proliferation of CNC cells ^(51)^, and to exert a positive effect on proliferation of dental mesenchymal cells ^(44)^. Our data show that FGF signaling rather than BMP and WNT signaling is highly active in human-mouse chimeric tooth germs where the differentiation duration of the mouse dental epithelium is largely slowed down (Fig. 2), which is consistent with the continuously high expression of FGFs, particularly FGF8, in human tooth germs, suggesting a potentially positive role of FGF signaling in regulating the tooth developmental tempo during odontogenesis in humans. Both our *ex vivo* lentivirus-mediated overexpression system and *in vivo* transgenic mouse model support that forcibly activated dental mesenchymal FGF8 signaling stimulates cell proliferation and inhibits cell apoptosis, thus prolonging tooth developmental pace in mice (Fig. 3-6). FGF8 appears to be a critical factor in regulating the tooth developmental pace during mammalian odontogenesis.

Functional redundancy complicates interpretation of FGF genetic studies. Phenotypes in various *Fgfr* knockout mice have demonstrated that different FGF receptors have both essential and redundant roles throughout development ^(49)^. During odontogenesis, FGF7 and FGF10 have been proven to act as surrogates of mesenchymal FGF3 ^(52)^, and FGF4, FGF8, as well as FGF9 have also been suggested functional redundancy and repetitive use in mouse tooth development ^(13)^. Our previous and current study show that several FGFs, especially FGF4, FGF8, and FGF9, have a broad and persistent expression in the whole tooth germs in humans but are restricted to the limited region of tooth germs in mice (Suppl. Table 1) ^(43)^. These data suggest that tooth developmental pace may be the product of synergistic effects of different FGFs in mammals. Actually, in addition to the fact that FGF8 is a key factor to control tooth developmental pace discovered in the current study, Fgf9 has also been reported to induce cell proliferation in cultured mouse dental mesenchyme *in vitro* ^(44)^ as well as in cervical loop development *in vivo* ^(53)^ and the *Fgf9* null mutant mice show a size-reduced labial cervical loop ^(53)^, and FGF4 stimulates cell proliferation in both the dental epithelium and mesenchyme as well ^(44,54)^, both suggesting their positive function on the regulation of tooth growth. We speculate that, in humans, several broadly, intensively, and persistently expressed FGF ligands may act in concert to confer growth capability (stimulation of proliferation and prevention of apoptosis) on tooth tissues to achieve longer development time-scale.

### FGF pathway may also participate in the regulation of tooth size

On the other hand, the developmental tempo and size of individual organs must be tightly coordinated to match the body size and larger-bodied species always keep slower developmental tempos to acquire larger organ sizes. Hippo signaling is the only admitted pathway that plays its best-known physiological function on the control of organ size in the mouse model for mammalian lineage ^(5-8)^, which mainly involves in the regulation of cell proliferation and apoptosis and thereby cell number ^(9)^. Activation or inhibition of YAP/TAZ, a transcriptional co-activator of Hippo pathway, leads to the adjustment of cell proliferation and apoptosis, thus affecting organ sizes and tissue homeostasis ^(5-8)^. However, in tooth development, suppression of YAP *ex vivo* at the cap stage only leads to ectopic cusp formation ^(55)^, while overexpression of YAP *in vivo* disrupts tooth morphogenesis with abnormal enamel knot as well as greatly widened dental lamina; teeth ultimately formed in YAP transgenic mice do not exhibit any growth delay or size enlargement ^(6)^. These data suggest that the renowned control function of Hippo pathway in mammalian organ size is not applicable for the tooth organ. In support of this notion, it has also been reported that size-control mechanisms differ across mammalian organs ^(11)^ and enhanced Hippo signaling induces organ enlargements in the stomach, heart, liver, and spleen, but not in lung, kidney, and intestine ^(56-58)^. Besides, IGF signaling is the other pathway that has been implied to have effects on organ size and *Igf2r* disrupted mice are born 25-35% bigger than wild-types, however, it has also been revealed that IGF signaling affects the size in some organs such as the cecum and colon but not the size of other organs such as the skin and stomach ^(10)^. In addition, it has indicated that tuning Bmp activity via enhanced epithelial Noggin activity modulates the size and number of the teeth in mice ^(24)^. Herein, we uncovered that natural broadly and persistently distributing pattern of FGF ligands, especially FGF8, in the human tooth germs may be also closely related to its tooth size.

### Dental mesenchymal FGF8 signal promotes cell cycle progression from G1-to S-stage via enhancing p38 and PI3K-Akt signaling to stimulate proliferation and prevent differentiation

Transduction of FGF signaling is mediated by multiple intracellular pathways, including P38, PI3K/Akt, Erk1/2, and JNK ^(49)^. In our previous study, the similar expression patterns of the indicators in Erk, P38, JNK, and PI3K-Akt pathways in mouse and human tooth germs suggests their conservative regulatory functions during mammalian odontogenesis ^(43)^. P38 has been reported to be involved in control of the cell cycle via the expression changes of P21, P27 and Ccnd3 in mouse skeletal muscle cells ^(59)^, Ccnd1 and c-Myc in human dental pulp stem cells ^(60)^, and Ccnd1 and P21 in rat mesenchymal stem cells ^(61)^, which is also significantly increased in our *Wnt1-Cre; R26R*^*Fgf8*^ mice (Fig. 7). The PI3K-Akt pathway is associated with a variety of cellular functions including cell growth, cell cycle regulation, and cell survival ^(62)^. The increased PI3K-Akt level regulates the proliferation and differentiation of neural progenitor cells by the activation of the cell cycle through the elevation of the Ccnd1 and cyclin E, two cyclins essential for the G1-S transition during organogenesis ^(63)^. Moreover, Akt protein is a central intracellular factor to transduce and integrate multiple extracellular growth factor signals, such as FGF, BMP, and GH/IGF, into a comprehensive signal network to supervise the processes of cell proliferation, growth, and survival ^(11)^. The increased activity of PI3K-Akt pathway caused by augmented FGF8 in the cranial neural crest cell (Fig. 7) further supports its positive function in cell proliferation regulated by cell cycle during tooth development. Erk has been reported to provide a switch-like transition between proliferation and differentiation of muscle progenitors ^(64)^, while JNK pathway has been elucidated to play crucial role in negative growth regulation and likely performs a central regulator in both local growth coordination and global growth coordination ^(4)^. These two pathways did not show any noticeable alteration in our *Fgf8* transgenic mice, which may attribute to their main function in control of negative growth or balancing cell proliferation and differentiation.

Cell growth and division are controlled by the cyclin depend kinases (CDKs) and their binding partners (cyclins) that regulate cell cycle progression through the regulation of the G1-S and G2-M transition. Many cell cycle regulators have been proved to control organ growth, which appears to be generally conserved in *Drosophila* and mammals ^(4)^. The G1-S phase transition is a rate-limiting step, a known checkpoint between cell growth and cell division, requiring cell cycle regulators to monitor and regulate the progression of the cell cycle ^(65)^. The stimulatory effects of cyclins can be offset by CDKs inhibitors (CDKIs) and the repression of CDKIs is also a prerequisite for G1-S transition. Thus merely elevating the level of cyclin D1 is not sufficient to induce cell cycle entry ^(66)^. Cyclin D1 (Ccnd1) is synthesized rapidly in the G1 phase and accumulates in the nucleus and degraded as the cell enters the S phase ^(45)^. P21, known as CDKI1, functions as a regulator of cell cycle progression in G1 and S phase and inhibits all cyclin/CDK complexes, which in turn results in cell cycle arrest and/or apoptosis ^(46)^. Enhanced dental mesenchymal FGF8 may promotes cell cycle progression through the regulation of the G1-S transition, which further stimulates cell proliferation and prevents cell differentiation and apoptosis, thereby resulting in prolonged tooth development. In line with this notion, we found that the amount of nuclear Ccnd1^+^ cells is dramatically increased while that of nuclear P21^+^ cells is markedly decreased in the dental mesenchyme at both the early and differentiation stage in *Wnt1-Cre;R26R*^*Fgf8*^ mice as compared to those in the control (Fig. 8), stating that forcedly enhanced FGF8 in the dental mesenchyme could elevate the level of Ccnd1 and meanwhile suppress the level of p21. Stimulation of Ccnd1 and repression of p21 both constitute the prerequisite for inducing cell cycle entry and thereby promoting the G1-S transition.

### FGF8 controls developmental pace in a non-cell-autonomous manner during early odontogenesis

Two latest studies have demonstrated that interspecies differences in developmental tempo are caused by cell-autonomous differences in biochemical reaction speeds instead of differences in signaling, the sequence of genes, or regulatory elements in *in vitro* human and mouse platforms ^(67,68)^. Moreover, in *Drosophila*, it is also reported that the *Minute* mutation causes a prolongation of developmental pace and decreased cell division rate in a cell-autonomous way via decreasing the expression of ribosomal proteins and thereby reducing the rate of translation and protein synthesis ^(69,70)^. Developmental allochrony regulated in a cell-autonomous manner is highlighted in these studies. However, our findings that the dental mesenchyme resorts to extrinsic signals such as FGF to dominate the developmental tempo as well via tissue interaction in a non-cell-autonomous manner during tooth development are inconsistent with these studies. Additionally, the fact that the oscillation phase shift between Notch and Wnt signaling is critical for the segmentation of presomitic mesoderm cells in multicellular organ development in 2D assays *ex vivo* ^(71)^ also indicates the pivotal role of secretory signal changes in the regulation of developmental pace. In *Drosophila*, organ growth are orchestrated not only by the growth rate regulated via signaling pathways but also by the growth duration regulated via hormonal signals and organ-autonomous processes controlled by patterning genes and physical force ^(4)^. Our heterogenous recombination results provide compelling evidence to support the contribution of extrinsic cues in the developmental tempo of teeth relying on tissue interaction during early development. Thus, we suggest that complicated regulatory networks involved in the developmental pace work both in a cell-autonomous and non-cell-autonomous way to orchestrate the organ morphogenesis, and, particularly, tissue interaction in a non-cell-autonomous way plays a major role during early organogenesis.

Tissue bioengineering based on the principle of developmental biology represents a promising approach for repair and regeneration of damaged tissues and organs in the future. However, unmatched development phase between humans and the model animals such as *Drosophila*, zebra fishes, and mice appear to be a major obstacle to acquire knowledge about organ growth control in humans. Uncovering of the molecular mechanism underlying organ growth control in humans is a prerequisite for the realization of replacement therapy in the future. Our study provides important FGF signaling information in tooth developmental process and enlightens our knowledge about the nature of FGF signaling in human tooth growth.

## Data Availability Statement

The data that support the findings of this study are available from the corresponding author upon reasonable request.

## Acknowledgments

This study was supported by the National Natural Science Foundation of China (81870739, 82001002, 81271102, 81771034) and the Natural Science Foundation of Fujian Province (2019J01281, 2020J01180).

