## Supplemental material for "FGF8-mediated signaling regulates tooth developmental pace during odontogenesis"

### Supplementary information

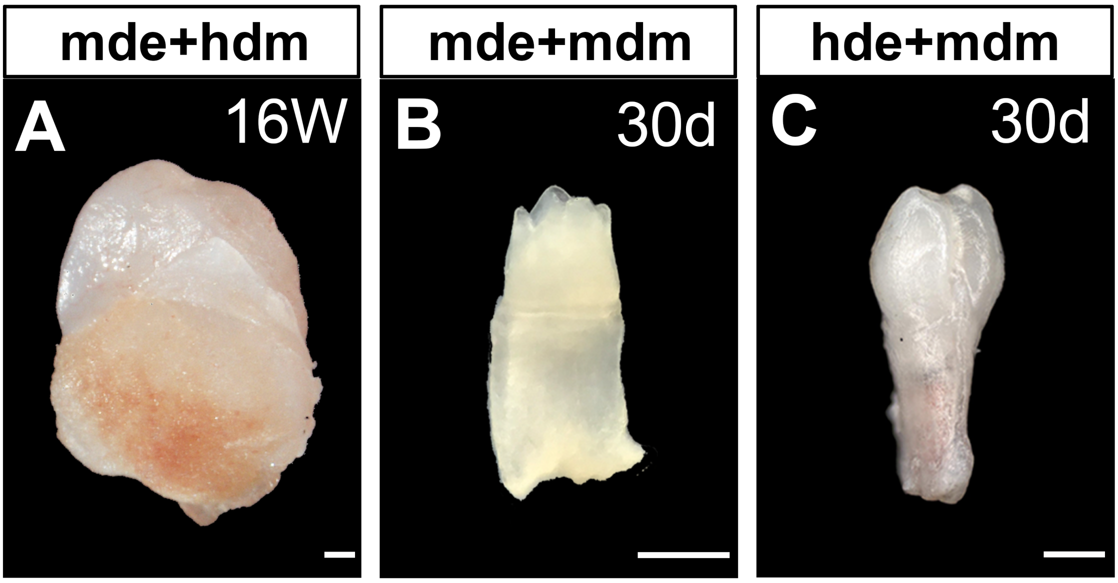


**Suppl. Fig. 1. Lateral view of different types of chimeric teeth after mineralization in subrenal culture.** A lateral view of tooth crowns in mde+hdm (**A**), mde+mdm (**B**), and hde+mdm (**C**) chimeras after 16 weeks, 30 days, and 30 days in subrenal culture, respectively. Scale bar = 500 µm.

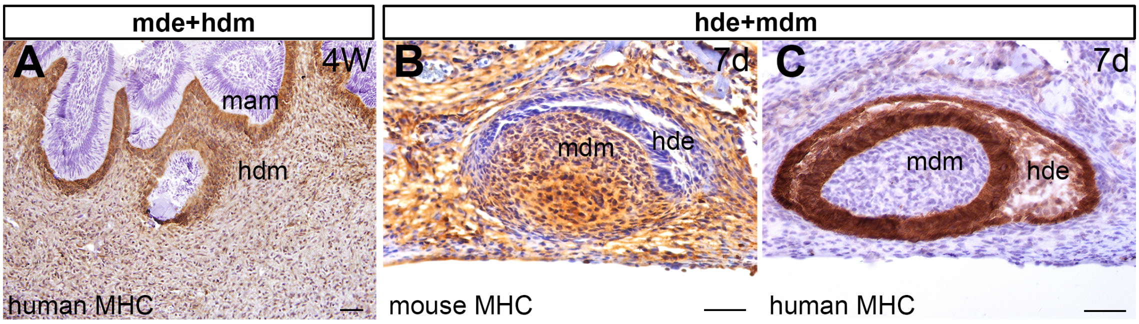


**Suppl. Fig. 2. No tissue contamination is confirmed by immunostaining of anti-MHC antibodies in the human-mouse heterogeneous recombination. A**: The mesenchymally exclusive expression of human MHC in recombinant of mouse dental epithelium and human dental mesenchyme after subrenal culture for 4 weeks. **B**: The mesenchymally exclusive expression of mouse MHC in the recombinant of human dental epithelium and mouse dental mesenchyme after subrenal culture for 7 days. **C**: The epithelially exclusive expression of human MHC in the recombinant of human dental epithelium and mouse dental mesenchyme after subrenal culture for 7 days. hdm, human bell-stage dental mesenchyme; mam, mouse ameloblasts; mdm, E13.5 mouse dental mesenchyme; hde, human bell-stage dental epithelium. Scale bar = 50 μm.

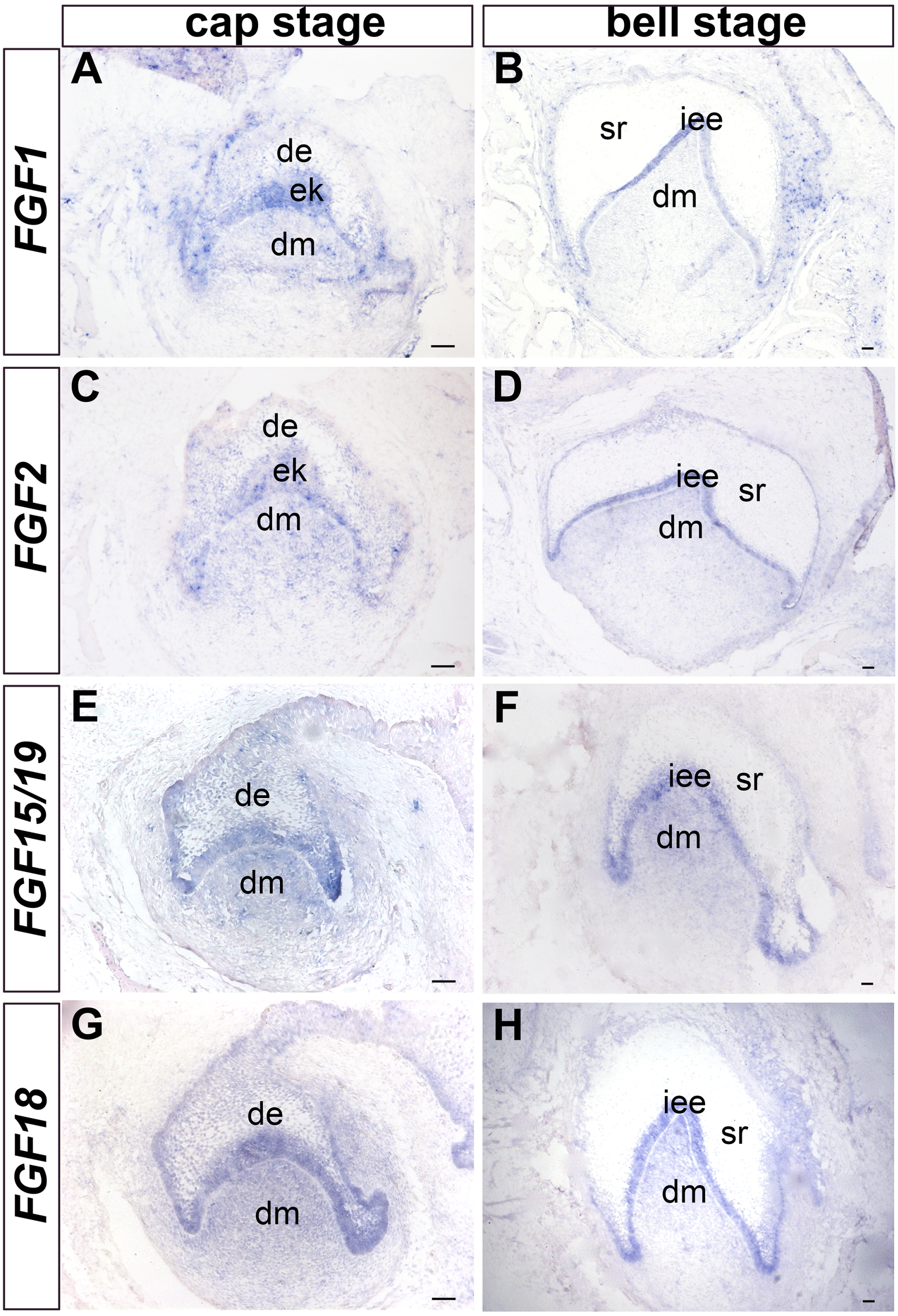


**Suppl. Fig. 3. Expression pattern of *FGF1*, *FGF2*, *FGF15/19*, and *FGF18* in human tooth germs at the cap and bell stages.** *In situ* hybridization shows the expression of *FGF1* (**A**,**B**), *FGF2* (**C**,**D**), *FGF15/19* (**E**,**F**), and *FGF18* (**G**,**H**) in human molar germs at the cap (**A**,**C**,**E**,**G**) and bell (**B**,**D**,**F**,**H**) stages. de, dental epithelium; dm, dental mesenchyme; ek, enamel knot; sr, stellate reticulum; iee, inner enamel epithelium. Scale bar = 50 μm.

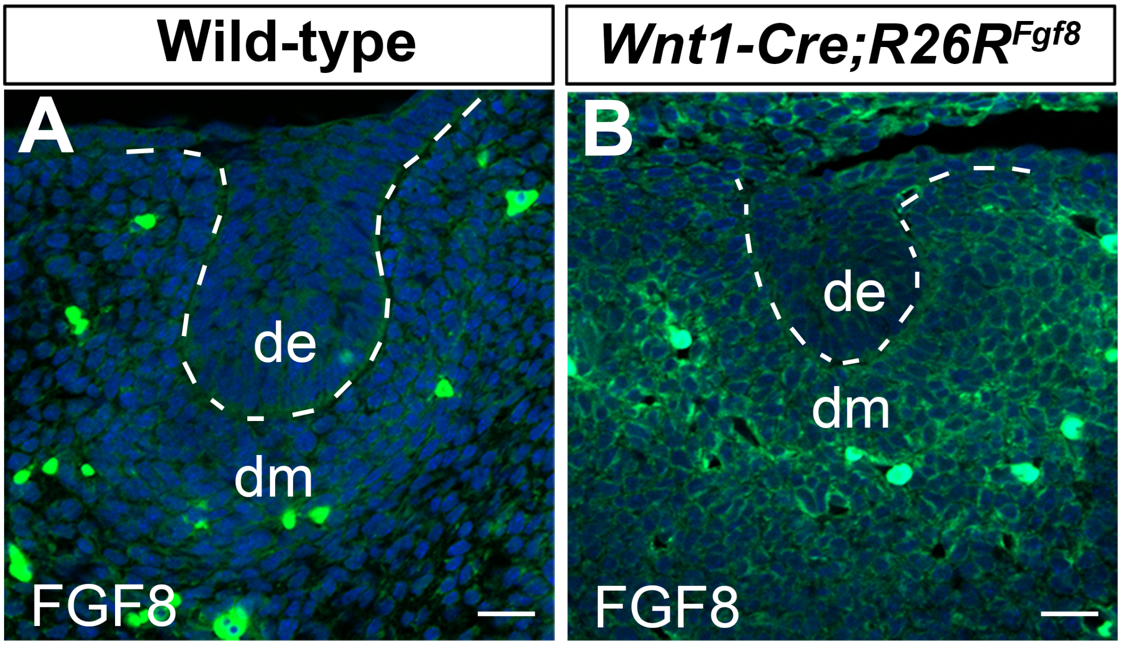


**Suppl. Fig. 4. *Fgf8* is ectopically expressed in E13.5 molar germs of *Wnt1-Cre;R26R$^{Fgf8}$* mice.** Immunostaining of FGF8 in E13.5 wild-type (**A**) and mutant (**B**) molar germs. de, dental epithelium; dm, dental mesenchyme. Scale bar: 50 µm.

**Suppl. Table 1. Comparison of the expression pattern of FGF ligands between human and mouse tooth germ at the cap and bell stages.**

| Gene | Mouse | | Human | |
|---|---|---|---|---|
| names | Cap stage | Bell stage | Cap stage | Bell stage |
| *Fgf1* | de | de; dm(slight) | de; dm(slight) | iee |
| *Fgf2* | de | sr | de; dm | iee |
| *Fgf3* | ek; dm | dm | de; dm | de |
| *Fgf4* | ek | ek | de; dm | de; dm |
| *Fgf5* | √ | × | √ | √ |
| *Fgf6* | × | × | × | × |
| *Fgf7* | × | × | de; dm(slight) | iee |
| ***Fgf8*** | × | × | de; dm | de; dm |
| *Fgf9* | ek | de | de; dm | de; dm |
| *Fgf10* | dm | dm | de (slight); dm(slight) | × |
| *Fgf15/19* | ek | dm | ek; dm | iee; dm |
| *Fgf16* | de; dm | de | × | × |
| *Fgf17* | de; dm | × | √ | √ |
| *Fgf18* | dm | × | de; dm | iee; dm |
| *Fgf20* | ek | ek | √ | × |
| *Fgf21* | ? | ? | × | × |
| *Fgf22* | ? | ? | × | × |
| *Fgf23* | ? | ? | √ | √ |

Abbreviations: de, dental epithelium; dm, dental mesenchyme; ek, enamel knot; iee, inner enamel epithelium; oee, outer enamel epithelium; sr, stellate reticulum. ?, unknown; ×, no expression; √, expressed but unknown whether in the dental mesenchyme or epithelium. Note that *FGF6*, *FGF16*, *FGF21*, and *FGF22* were undetectable in human tooth germs. Although *FGF1*, *FGF2*, *FGF3*, *FGF7*, *FGF10*, and *FGF20* were detected in human tooth germs at the cap stage, they were either undetected or only expressed in the dental epithelium at the bell stage, indicating that these genes were not continuously expressed in the dental mesenchyme of human tooth germs. Despite the fact that *FGF5*, *FGF15/19*, *FGF17*, and *FGF18* were detected in human tooth germs (including dental mesenchyme) at the cap and bell stages, they were also detected in mouse dental mesenchyme either at the cap stage or the bell stage, suggesting that these genes were not specifically expressed in human dental mesenchyme. In addition, *Fgf23* is excluded because it is expressed not only in the dental epithelium but also in the dental papilla at the differentiation stage (1), which is consistent with its expression pattern in human tooth germs (2). By comparison, *FGF4*, *FGF8*, and *FGF9* exhibit continuous and broad expression in human tooth germs but strictly restricted in the mouse tooth germ.

**Suppl. Table 2. Primer sequences for each gene using in RT-PCR.**

| **Gene** | **Forward Primer** (5'-3') | **Reverse Primer** (5'-3') | **Size** (bp) |
|---|---|---|---|
| *FGF1* | GAGCCTGAATTTGTAAGCAAC | TGCCATCCTTCTAAGTTACCC | 470 |
| *FGF2* | CGGTCAAGGAAATACACCAGT | AGAACTTCCAGCAGTTTACACA | 645 |
| *FGF5* | TGCCAAGATTCAAGCAGTCG | CCCTGTTATTTAACTTTCCGAA | 456 |
| *FGF6* | AATTCTGACCACGTGCCTGA | TAAGCCTTCTTTTGTGGGTCCT | 680 |
| *FGF16* | CGTGCCCTTAGCTGACTCC | ATCTTTGTTCAGGGCCACGTA | 433 |
| *FGF17* | CGCCTGCTGCCCAACCTCA | GCCATGAACCAGCCCTCGT | 443 |
| *FGF18* | ATGACAAAAGACTCACGCAAA | AGCTGGTTTTAAAATATTCCCT | 504 |
| *FGF15/19* | GAACAGTCCTGAGTCCACGTT | CATGCCTGCTCCAGTCAGT | 476 |
| *FGF20* | GGACCACAGCCTCTTCGGTA | GCCATCTCTTGGAGTTCCGTCT | 265 |
| *FGF21* | TCTGGCACCAATTCTAAACCAC | GATTTGAATAACTCCCGGCTT | 353 |
| *FGF22* | GCGCCTCTTCTCCTCCACTCACT | GCCGTTCTCTTCGATGCGCTCCC | 241 |
| *FGF23* | TTAAATGAAGCCTTACCCCAT | AAATACTGCCACATGACGAG | 523 |
| *Fgf5* | GGAAGCGGCTCGGAACATAG | AGGAAAGTTCCGGTTGCTCG | Transcript variant-1 475<br>Transcript variant-2 371 |
